# Chickpea (*Cicer arietinum* L.) root system architecture adaptation to initial soil moisture improves seed development in dry-down conditions

**DOI:** 10.1101/2020.09.24.311753

**Authors:** Thibaut Bontpart, Ingrid Robertson, Valerio Giuffrida, Cristobal Concha, Livia C. T. Scorza, Alistair J. McCormick, Asnake Fikre, Sotirios A. Tsaftaris, Peter Doerner

## Abstract

Soil water deficit (WD) impacts vascular plant phenology, morpho-physiology, and reproduction. Chickpea, which is mainly grown in semi-arid areas, is a good model plant to dissect mechanisms involved in drought resistance.

We used a rhizobox-based phenotyping system to simultaneously and non-destructively characterise root system architecture (RSA) dynamics and water use (WU) patterns. We compared the drought-adaptive strategies of ‘Teketay’ to the drought-sensitive genotype ICC 1882 in high and low initial soil moisture without subsequent irrigation.

WD restricted vegetative and reproductive organ biomass for both genotypes. Teketay displayed greater adaptability for RSA dynamics and WU patterns and revealed different drought adaptive strategies depending on initial soil moisture: escape when high, postponement when low. These strategies were manifested in distinct RSA dynamics: in low initial soil moisture, its reduced root growth at the end of the vegetative phase was followed by increased root growth in deeper, wetter soil strata, which facilitated timely WU for seed development and produced better-developed seeds.

We demonstrate that RSA adaptation to initial soil moisture is one mechanism by which plants can tolerate WD conditions and ensure reproduction by producing well-developed seeds. Our approach will help in identifying the genetic basis for large plasticity of RSA dynamics which enhances the resilience with which crops can optimally adapt to various drought scenarios.

**Highlight:** Root system architecture and water use patterns change dynamically for distinct drought adaptation strategies in chickpea.

## Introduction

Water deficit (WD), or drought, is a major abiotic factor limiting plant growth and reproduction. Plant responses to drought are complex; they depend on its duration, frequency, intensity, the plant species/genotypes, as well as the timing of drought occurrence during the plant’s life cycle (Bodner *et al*., 2015; Tardieu, 2012).

Chickpea (*Cicer arietinum* L.) is globally one of the most highly produced grain legumes (FAOSTAT: http://www.fao.org/faostat/en/#data/QC). Mainly grown under rainfed conditions in (semi-) arid areas, the seeds are planted at the end of the rainy season to avoid water-logging (Mohammed *et al*., 2017). The majority of chickpea is produced by smallholders who lack access to water for irrigation to ensure optimal growth conditions (Hanjra *et al*., 2009). As a result, the plant grows on varying levels of residual soil moisture which decrease over the growing season. This dry-down scenario can lead to flower and pod abortion and ultimately, decreases grain yield (Fang *et al*., 2010; Leport *et al*., 1999; Pang *et al*., 2017a; Pang *et al*., 2017b). Chickpea performance depends on environmental constraints (*e.g*. rainfall regime, evaporative demand, soil type), but also on its genetic background, which drives its response to drought. In the tropics and sub-tropics, water resources and grain legume yield are predicted to decrease due to climate change (Andrews and Hodge, 2010). The selection of genotypes best adapted to local environmental constraints will be crucial for resilient, sustainable agriculture in a context of climate change (Singh *et al*., 2014).

In experiments, chickpea responses to drought are commonly studied with experimental designs that start with growth in well-watered conditions, followed by cessation of irrigation to simulate ‘terminal drought’ at mid-vegetative stage (Zaman-Allah *et al*., 2011b), early podding stage (Pang *et al*., 2017a; Pang *et al*., 2017b), or at different podding stages (Leport *et al*., 2006). In the field, chickpea performance was evaluated comparing optimal irrigation with rainfed (drought stress) conditions (Purushothaman *et al*., 2017a, b). However, the former conditions do not recapitulate the conditions in fields planted after cessation of seasonal rains.

Against a backdrop of climate change, the combination of lower rainfall and elevated temperatures will accelerate evaporation, leading to lower soil moisture at sowing and during the growth period. However, very few studies describe the impact of initial soil moisture on chickpea performance (*e.g*. Hosseini *et al*., 2009) and early WD was only tested transiently (Fang *et al*., 2011). Initial low soil moisture restricts shoot traits, such as height and biomass (Hosseini *et al*., 2009), but no information on chickpea root and reproductive traits have been reported.

In dry-down scenarios, managing available soil water use is critical for plants to optimize their reproduction. In the context of the drought resistance framework, a distinction is made between annual plants that *escape, postpone* or *tolerate* drought (Berger *et al*., 2016; Levitt, 1980). In chickpea, all three strategies to WD resistance have been observed (Berger *et al*., 2016), but their relevance is still controversial: *rapid-cycling* genotypes reach their reproductive stage quickly and are consequently less affected, as they can escape terminal drought (Kumar and Abbo, 2001). However, their short life cycle restricts growth and imposes a ceiling on yield (Krishnamurthy *et al*., 2010). By contrast, genotypes that *postpone* drought effects have specific water use (WU) patterns that facilitate coping with terminal drought: lower WU during their vegetative phase, then higher during the reproductive phase (Zaman-Allah *et al*., 2011b). The capacity to retain soil water (low transpiration rate per leaf area, TR) when it is abundant during the vegetative stage, even with a high vapour pressure deficit, was highlighted as a relevant trait to alleviate terminal drought since it allows to economise water for reproduction (Zaman-Allah *et al*., 2011a). Low TR genotypes exhibit early vigour (growth) and low root hydraulic conductivity (Sivasakthi *et al*., 2017). Low TR in early stages of water stress followed by high TR during the reproductive phase was suggested as the best water management strategy with respect to reproduction and yield (Pang *et al*., 2017a).

Field experiments have shown that high root length density (per volume unit) and root dry weight, both in deep soil strata, had a positive influence on chickpea yield in drought conditions (Kashiwagi *et al*., 2006; Purushothaman *et al*., 2017b). A recent field study comparing 12 chickpea genotypes has revealed that in drought conditions, (*i*) water uptake was positively linked to root length density in all soil strata, (*ii*) water uptake in deep soil strata (90-120 cm) ensured the best drought adaptation (Purushothaman *et al*., 2017a). Hence, investment in a dense, deep root system during the reproductive phase is a viable strategy to cope with dry-down. However, due to the technical difficulties of simultaneously and accurately measuring whole root system growth and whole-plant WU dynamics, the role of root system dynamics to secure optimal reproduction in WD conditions has not been well-studied.

Soil-filled phenotyping systems such as rhizoboxes are useful to study root system architecture (RSA) dynamics in soil-grown plants (Bodner *et al*., 2015; Bontpart *et al*., 2020; Nagel *et al*., 2012). RSA reflects, through a set of biologically relevant parameters, the spatiotemporal progression of root system development. Specifically, total root length and root area (RA) represent plant belowground biomass investment, whereas the convex hull area (CHA) is a proxy for the explored soil volume. Root system solidity (RA/CHA) reflects how intensively roots develop within the explored soil area. Root system depth and root distribution in progressively deeper soil strata are particularly relevant parameters to study root plasticity when water is more abundant at depth. The impact of initial soil moisture was tested during the early vegetative phase in rhizoboxes on sibling pre-breeding lines in wheat (Nagel *et al*., 2015), and maize hybrids from different geographic origins (Avramova *et al*., 2016). Studies of RSA dynamics hold great promise to dissect plant WU and growth strategies to study their contribution to yield maintenance.

To investigate that question, we used a large rhizobox system developed to follow RSA (Bontpart *et al*., 2020) and optimised it to study WU pattern non-invasively until the chickpea pod-filling stage. The recently released chickpea genotype (Teketay) and the drought sensitive genotype (ICC 1882) were grown in high (control, CT) and low (water deficit, WD) initial soil moistures. In addition to analysing RSA and WU kinetics, a set of shoot and yield-related traits were also quantified to determine (*i*) the impact of initial soil moisture on morpho-physiological traits and reproduction, (*ii*) the drought adaptive strategies developed by both genotypes in each soil moisture condition.

## Materials and methods

### Plant material and growth conditions

Seed of ICC 1882 and Teketay (accession number ICCV-00104) genotypes of desi chickpea (*Cicer arietinum* L.) were provided by the International Crops Research Institute for the Semi-Arid Tropics (ICRISAT) and the Debre Zeit Agricultural Research Center (DZARC), respectively.

Experiments were conducted in a greenhouse at the King’s Building campus (Edinburgh, UK, 55°55’14.9”N, 3°10’09.9”W). Twenty-four plants (5-7 replicates per genotype in both soil moisture conditions) were grown in the rhizobox system described by Bontpart *et al*. (2020).

John Innes seed compost (Levington, UK) was prepared so that initial soil water content (SWC; w:w) was 62% and 40% for CT and WD conditions, respectively. Before soil loading, the volumetric water content (VWC) was measured using an ECH2O EC-5 probe connected to Em5b Analog Data Logger (Decagon Devices), and adjusted to ~0.36 and 0.2 cm^3^ H_2_O/cm^3^, for CT and WD conditions, respectively.

The top of rhizoboxes was sealed with PVC electrical tape to suppress soil evaporation. Unplanted rhizoboxes sealed with PVC electrical tape were repeatedly weighed to ensure this largely prevented water evaporation. Average water loss from those mock rhizoboxes was 1.6 and 1.3 g d^-1^ for CT and WD, respectively. No extra water was added during the experiment.

Rhizoboxes were randomly distributed between two supports. Plants were separated by 12 cm within the same row, which is similar to plant spacing in the field (Gaur *et al*., 2010). Chickpeas were grown under daylight supplemented with ~200 μmol m^-2^ s^-1^ at plant height, provided by HPS lamps from 6 am to 6 pm if external light dropped below a threshold of ~1000 μmol m^-2^ s^-1^. Air temperature and relative humidity was recorded with a USB-502 data logger (Measurement Computing, USA) placed at canopy level (Supplementary Fig. S1 at *JXB* online). The average night and day temperature were 18.7 ± 0.1 °C and 26.8 ± 0.1 °C, respectively. The average night and day relative air humidity were 63.9 ± 0.4% and 46.7 ± 0.4%, respectively.

### Plant measurements

#### Root traits

Images of each rhizobox were taken twice a week for seven weeks, starting from 6 until 52 days after sowing (das), using an imaging station equipped with an array of Phenotiki imaging devices (Bontpart *et al*., 2020; Minervini *et al*., 2017). Image processing and RSA parameters were computed as described in Bontpart *et al*. (2020). Root area (RA) was calculated separately for 10 horizontal soil strata of 15 cm each covering the whole rhizobox. For each layer, the difference in root area (ΔRA) for the same genotype between soil moisture conditions, were calculated as RA_CT_ – RA_WD_; and ΔRA between genotypes grown in the same soil moisture condition as RA_Teketay_ – RA_ICC 1882_. To determine the length of the root system each seed pod potentially benefits from with respect to water extraction, we calculated the ratio between pod number and root system length. At 53 das, the root system was washed, blotted dry, and weighed to evaluate its fresh mass (FM), then oven-dried (72 h at 70°C) for dry mass (DM).

#### Shoot traits

Plant height was estimated by measurement of the distance between the point of emergence from the soil and the meristem of the main stem. At 52 das, shoots were excised and partitioned into leaves, stem, flowers, and pods to weigh FM. Shoot material was oven-dried (72 h at 70 °C) then weighed for DM. Biomass investment in each organ, related to total plant biomass, and the root/shoot ratio (R/S), were calculated on DM basis and expressed as percentage. After drying, pods were crushed to extract, count and weigh each seed. The final seed set was calculated as the percentage of pods containing at least one seed. The mean seed mass (0.287 g and 0.228 g for Teketay and ICC 1882, respectively), “optimal seed mass”, was calculated by weighing 100 seeds from the master stock used for the experiment, which had been propagated in optimal conditions. The developmental index was determined for each seed, by comparing its mass to the optimal seed mass of the corresponding genotype. The average seed development index was calculated for each plant. Seeds were divided into four classes according to their development index: > 75%, 75-50%, 50-25% and < 25%. The harvest index was determined as the ratio between seed DM and whole-shoot DM.

#### Reproductive development

Flowering time (FT) was determined as das until the first flower opened. To track reproductive development and success, the number of reproductive organs (sum of open flowers and pods) was counted. The number of pods was counted from the day the first visible pod(s) (few mm long) appeared. The pod set was calculated as the ratio of pods per reproductive organs.

#### Water use

Just after image acquisition, each rhizobox was weighed on a platform scale (IFB 30K-3, Kern, 0.001 kg readability) to evaluate plant WU as the rhizobox mass loss. The WU rate (WUR) was calculated between two consecutive weighings as follows:

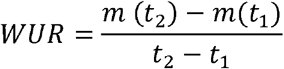

where *m*(*t*) is the rhizobox mass at time *t*. The total WU (TWU) corresponds to the rhizobox mass difference between the first and the last weighing. The water use efficiency (WUE) is the ratio whole-shoot DM/TWU. Final transpiration rate per leaf mass (TR_mass_) was estimated as the difference in WUR between 52 and 48 das, divided by leaf DM at harvest. To estimate how well pods are irrigated, we calculated the ratio between WUR and pod number (g H_2_O d^- 1^ pod^-1^).

#### Leaf gas exchange

Gas exchange was measured on ICC 1882 chickpeas with a LI-COR 6400-40 Leaf Chamber Fluorometer connected to a LI-6400XT Portable Photosynthesis System (LICOR), using the same parameters and leaves as described by Pang and colleagues (2017a; 2017b). Leaves used for gas exchange measurements were detached and scanned. Leaf area was determined with the ImageJ wand tool after conversion to 8-bit colour and threshold adjustment with default parameters. Net photosynthetic CO_2_ assimilation rate and stomatal conductance were reported per unit leaf area (*A* and *g*s, respectively). We measured them when the first flower opened, to examine the impact of initial soil moisture levels on chickpea leaf gas exchange (Supplementary Fig. S2). Both traits were significantly (*p* < 0.05) lower in WD conditions.

#### Soil water content

At the end of the experiment and just before root harvest, soil samples (~10-15 g) were collected at ~7.5, 37.5, 82.5, and 135 cm depth from planted rhizoboxes. Soil samples from unplanted rhizoboxes were collected in the middle of the experiment (26 DAS) and at harvest (Supplementary Fig. S3). Fresh soil samples were weighed, then oven-dried (72 h at 70 °C), to determine soil water content (SWC) expressed on a gravimetric basis:

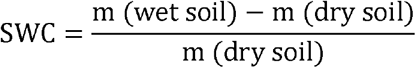

The SWC of unplanted rhizoboxes were subtracted from the SWC of planted rhizoboxes of the corresponding water regime to calculate ΔSWC reflecting water consumption.

#### Statistics

The effects of soil moisture and genotype on trait kinetics were analysed independently using one-way ANOVA. All statistics analyses were performed with Infostat 2019.

## Results

### Low initial soil moisture restricts growth and water use of both genotypes

We analysed the impact of initial soil moisture on (*i*) growth (shoot height, RSA), physiological (Water Use, WU) and reproductive traits (reproductive organs, pod number, pod set), (*ii*) the kinetics of changes in pod number per root length, and Water Use Rate (WUR) per pod, and (*iii*) parameters calculated after harvest (Water Use Efficiency, WUE; biomass distribution; final Transpiration Rate per leaf mass unit, TR_mass_; seed development index; harvest index) to establish whether the parallel analysis of these parameters provided insights into drought response strategies.

Shoot height was used as a proxy for shoot growth. It was significantly reduced by WD from 10 and 17 das for Teketay and ICC 1882, respectively (Fig. 1). Shoot growth restriction imposed by WD is paralleled in root growth: total root length is significantly reduced from 13 das onwards for both genotypes (Fig. 2A). Teketay convex hull area (CHA) and root depth were markedly reduced by WD from 17 das onwards (Fig. 2B, C). ICC 1882 CHA was significantly lower in WD conditions at 17 das (*p* < 0.05), then from 24 das onwards, while its root depth was significantly reduced from 10 das onwards. Root system solidity of both genotypes was significantly lower in WD conditions from 20 and 24 das for Teketay and ICC 1882, respectively (Fig. 2D). Root growth rate (RGR) was significantly impacted by WD except in very early (at 10 das for both) and late stages (45-52 das for Teketay, 48-52 das for ICC 1882) when RGR also declines in CT conditions (Fig. 2E, F).

**Figure 1.**
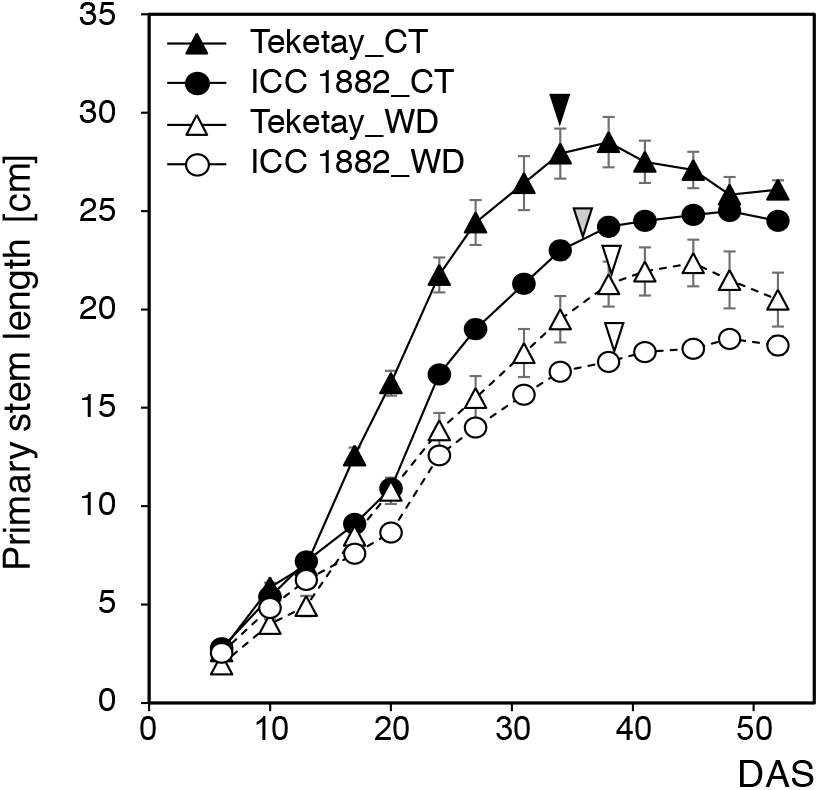
Time course of shoot height of the chickpea genotypes Teketay and ICC 1882 in control (CT) and water deficit (WD) conditions. Data are mean values (±SE) of five to seven plants. The black, grey and white arrows mark the average flowering time for Teketay in CT, ICC 1882 in CT and both genotypes in WD conditions, respectively.

**Figure 2.**
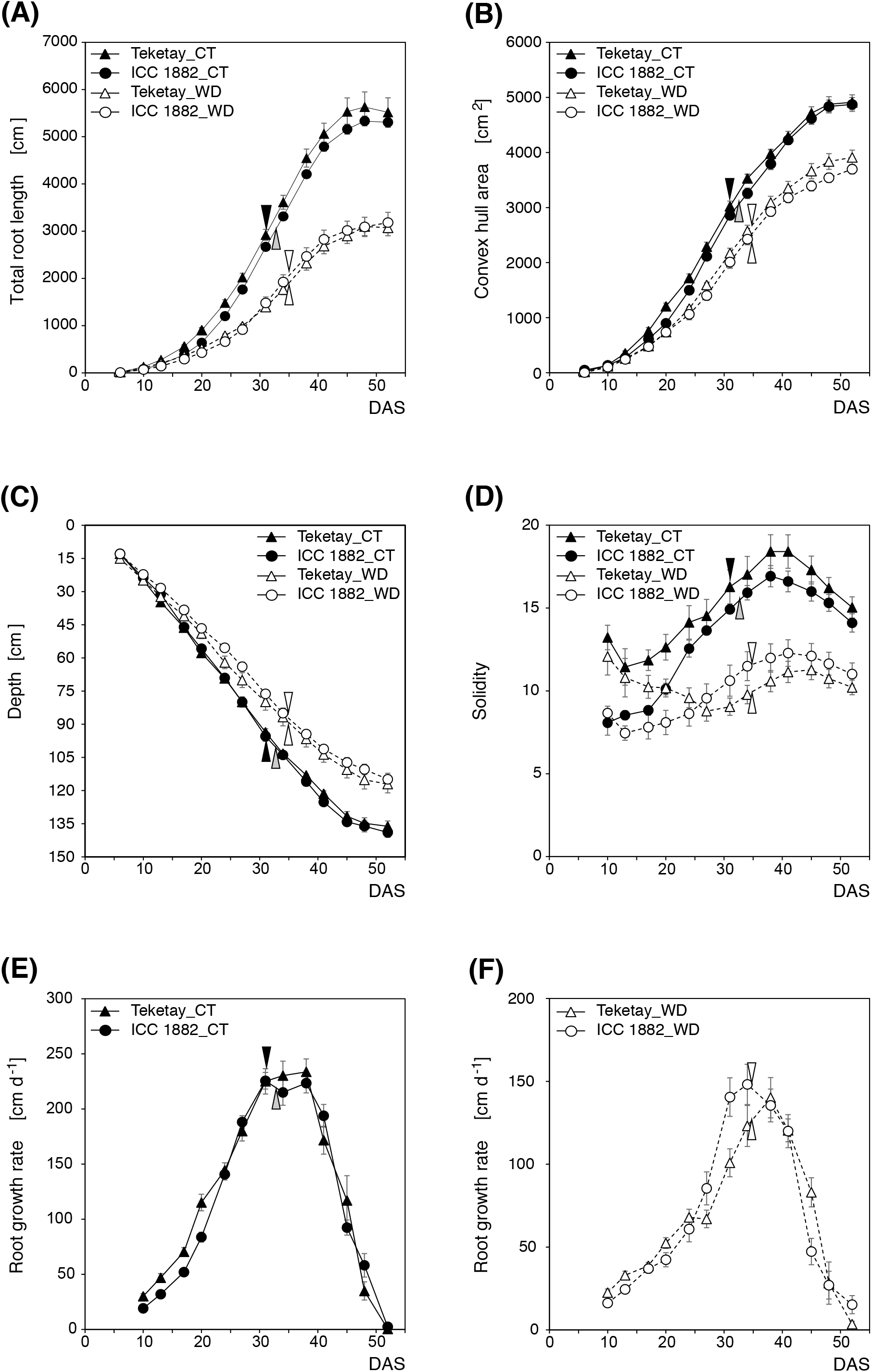
Root system architecture parameters of the chickpea genotypes Teketay and ICC 1882 in control (CT) and water deficit (WD) conditions. Total root length (a), convex hull area (b), depth (c), solidity (d), root growth rate in CT conditions (e) and WD conditions (f). Data are mean values (±SE) of five to seven plants. The black, grey and white arrows mark the average flowering time for Teketay in CT, ICC 1882 in CT and both genotypes in WD conditions.

Overall, WD reduced all RSA parameters (length, CHA, depth, area, solidity, RGR) from the beginning of the vegetative phase. At the end of the experiment, contrasting root system phenotypes illustrate WD impact (Supplementary Fig. S4), and their computed root area and length are well correlated with the harvested root fresh and dry mass (Supplementary Fig. S5). ΔRA_CT-WD_ was always positive, indicating higher root growth in CT conditions in all soil strata for both genotypes (Supplementary Fig. S6), with no evidence of a global stimulation of root growth induced by WD for water “foraging”. This is clearly visible when comparing the RA spatiotemporal evolution between CT and WD conditions for Teketay (Supplementary Fig. S7A, C, respectively), and ICC 1882 (Supplementary Fig. S7B, D, respectively). Along with reduced root system development in WD, cumulative WU was significantly reduced by WD for both genotypes (except at 10 das for ICC 1882, Fig. 3A), resulting in a total decrease of WU by ~60% (Table 1). WUR was significantly reduced for both genotypes (Fig. 3B). In leaves, WD significantly increased final TR_mass_ for Teketay (*p* < 0.05, Table 1).

**Figure 3.**
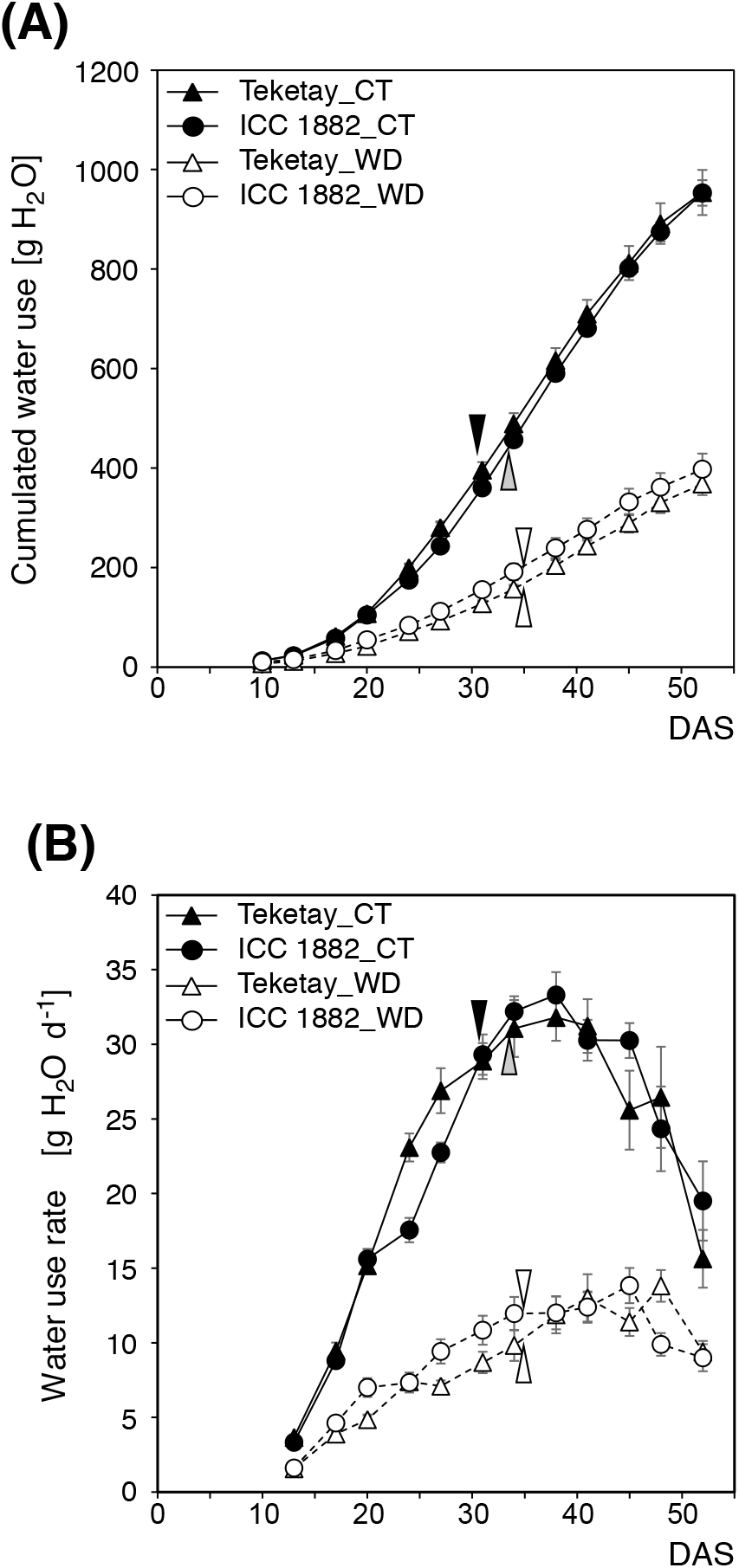
Water use of the chickpea genotypes Teketay and ICC 1882 in control (CT) and water deficit (WD) conditions. Cumulated water use (a) and water use rate (b). Data are mean values (±SE) of five to seven plants. The black, grey and white arrows mark the average flowering time for Teketay in CT, ICC 1882 in CT and both genotypes in WD conditions, respectively.

**Table 1.**
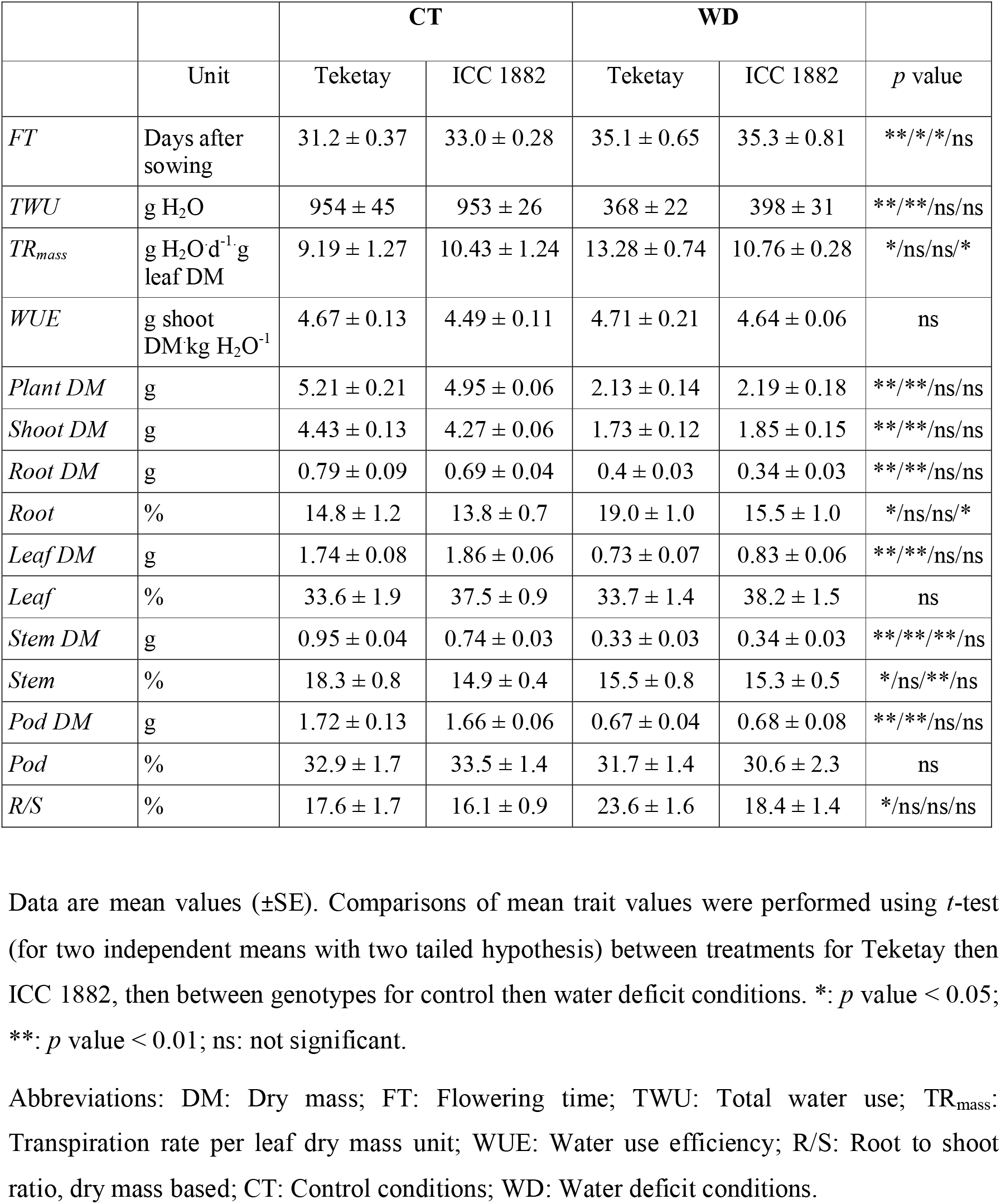
Flowering time and morpho-physiological traits of the chickpea genotypes Teketay and ICC 1882 in control (CT) and water deficit (WD) conditions grown in rhizoboxes.

For both genotypes, remaining soil moisture (SWC) was more abundant in WD conditions for all depths except at 135 cm where SWC values were very close to those in CT conditions (Fig. 4A). This indicates that remaining soil moisture in CT conditions is lower despite a higher initial SWC. The ΔSWC, which reflects plant water consumption, was significantly higher in CT conditions for both genotypes at all rhizobox depths (Fig. 4B).

**Figure 4.**
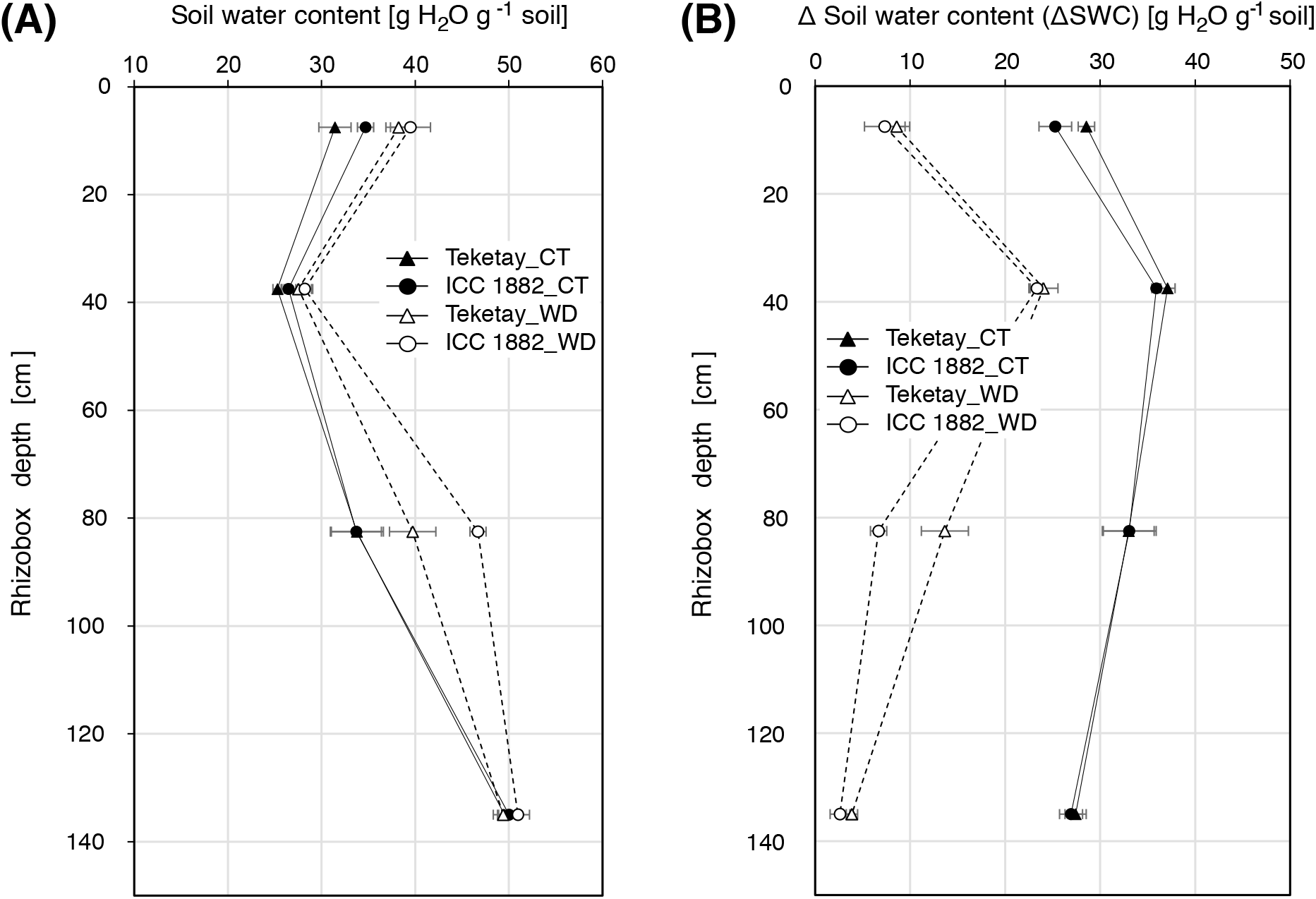
Soil water content (SWC) at different soil layers in rhizoboxes used to grow the chickpea genotypes Teketay and ICC 1882 in control (CT) and water deficit (WD) conditions. SWC (a) and SWC normalised (ΔSWC) to SWC measured in unplanted, soil-only rhizoboxes (b). Data are mean values (±SE) of five rhizoboxes.

### Low initial soil moisture alters Teketay resource allocation

At harvest, chickpea shoot material was separated into pods, leaves and stems to quantify organ biomass and biomass distribution (Table 1). WD significantly decreased plant DM (by −59.2 and −55.9% for Teketay and ICC 1882, respectively), shoot DM (by −61 and −56.7%), leaf DM (by −58.2 and −55.4%), filled pod DM (by −61.3 and −59.1%) and root DM (by −48.9 and 50.7%) in similar proportions. WD impacted Teketay more than ICC 1882 stem DM: by −65.3 and −54.6%, respectively. Harvest index was not significantly affected by WD, but its reduction was lower for Teketay compared to ICC 1882 (−6.1 and −16.2%, respectively). Teketay biomass allocation to roots was stimulated by WD while allocation to the stem was reduced, resulting in higher R/S in WD condition (*p* < 0.05 for all, Table 1). By contrast, WD did not impact biomass distribution for ICC 1882, and WD did not significantly affect WUE for both genotypes.

### Water deficit delays reproductive development and reduces pod-set

We used the opening of the first flower(s) to mark the transition between vegetative and reproductive phases and used this timepoint to calculate Flowering Time, FT. Compared to CT conditions, FT in Teketay and ICC 1882 growing in WD conditions was significantly delayed by 4 and 2.3 days on average, respectively (Table 1). At harvest (52 das), WD significantly decreased the number of reproductive organs (sum of flowers and pods) by 74.1% and 48.1% for Teketay and ICC 1882, respectively (Fig. 5A). For both genotypes, WD conditions reduced the number of pods significantly at all time points (Fig. 5B). Pod set, i.e. the percentage of flowers producing pods, was significantly lower for Teketay in WD conditions at 38 das, but higher from 45 das (*p* < 0.05 only at 45 das) compared to Teketay in CT conditions (Fig. 5C).

**Figure 5.**
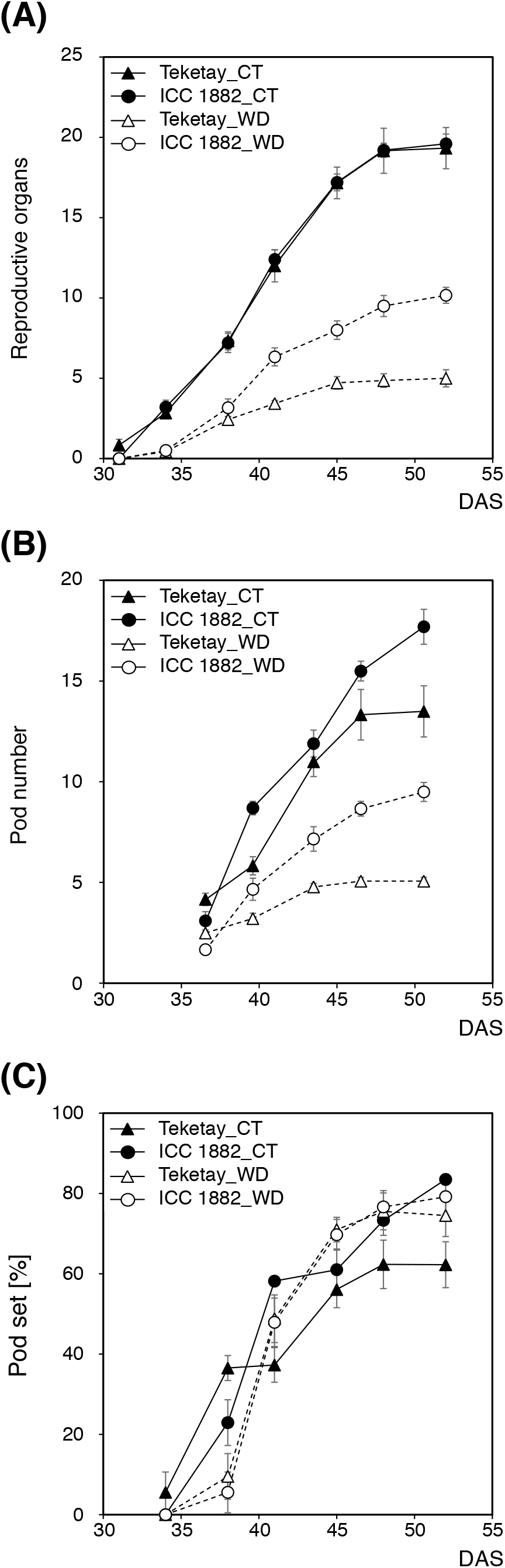
Time course of reproductive organ development in the chickpea genotypes Teketay and ICC 1882 in control (CT) and water deficit (WD) conditions. Number of reproductive organs (a), number of pods (b), pod set percentage (c). Data are mean values (±SE) of five to seven plants.

For comparison, we assumed an equal water extraction capacity per unit length of root (Fig. 6). Thus, the ratio between pod number and unit root length reflects the number of pods this normalised water extraction capacity supports for seed development. Overall, it tends to increase over time as the number of pods increases faster than the root system during the reproductive phase. It was significantly reduced by WD from 41 das for Teketay only (Fig. 6A). As a result, Teketay WUR per pod was significantly higher in WD conditions from 45 DAS but similar throughout the reproductive phase for ICC 1882 (Fig. 6B).

**Figure 6.**
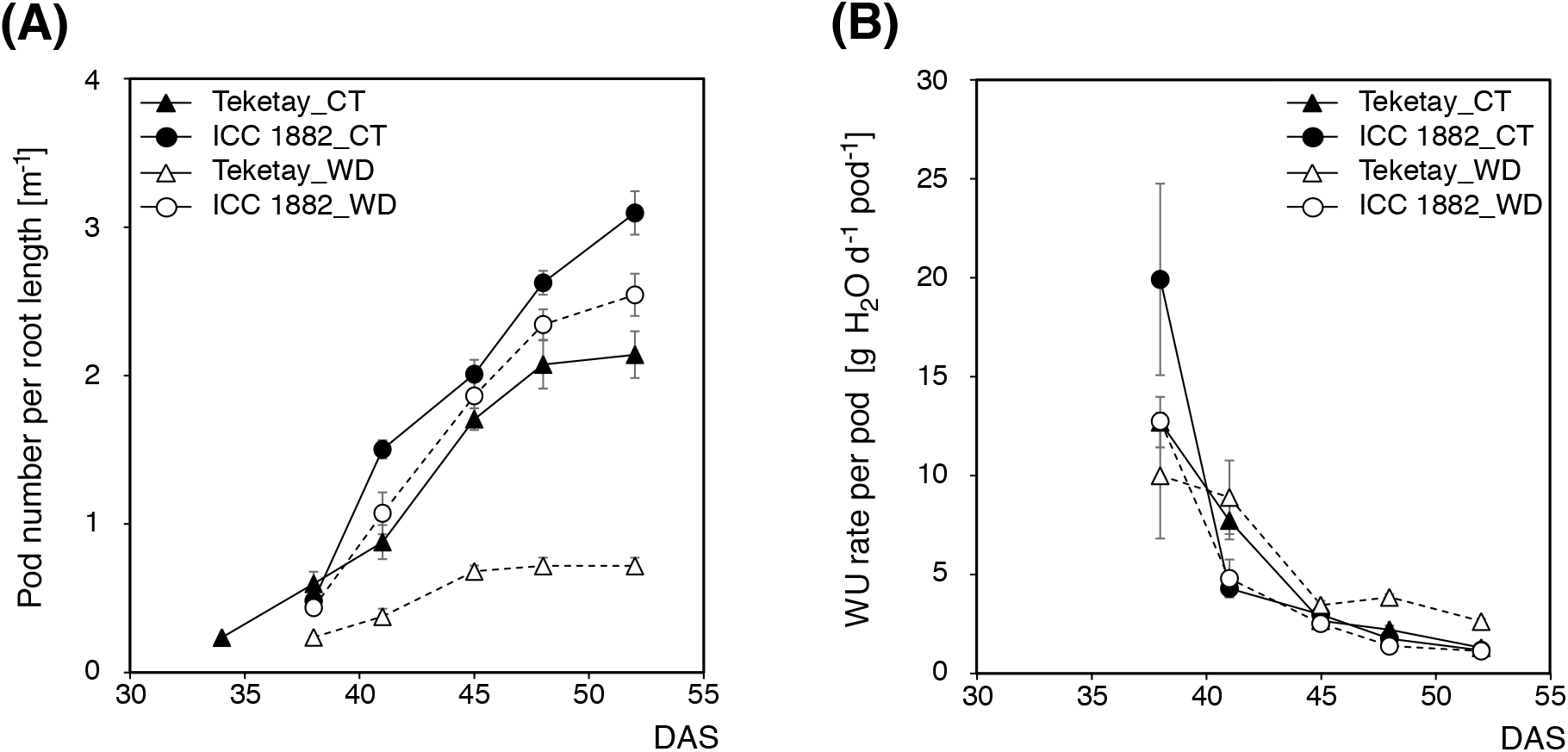
Root system length available for pod water supply in the chickpea genotypes Teketay and ICC 1882 in control (CT) and water deficit (WD) conditions. Number of pods per unit root system length (a) and water use rate per pod (b). Data are mean values (±SE) of five to seven plants.

### Water deficit reduces seed number and maturation

WD significantly decreased total seed mass by a similar degree for Teketay (−64%) and ICC 1882 (−62.5%, Table 2). WD also significantly reduced seed number, but more for Teketay (−58.5%) than for ICC 1882 (−46.9%). Strikingly, WD restricted average seed mass and development index for ICC 1882 (*p* < 0.05) but not for Teketay. Seed set, i.e. the percentage of pods containing at least one seed, in WD conditions was significantly higher for Teketay (100% versus 78.4% in CT condition), but not significantly different for ICC 1882 (83% versus 74.8% in CT condition). Only some Teketay plants grown in CT conditions produced seeds that reached the highest class of seed development index (> 75%) (Fig. 7). Seed distribution was not significantly impacted by WD for the other classes of seed development index.

**Figure 7.**
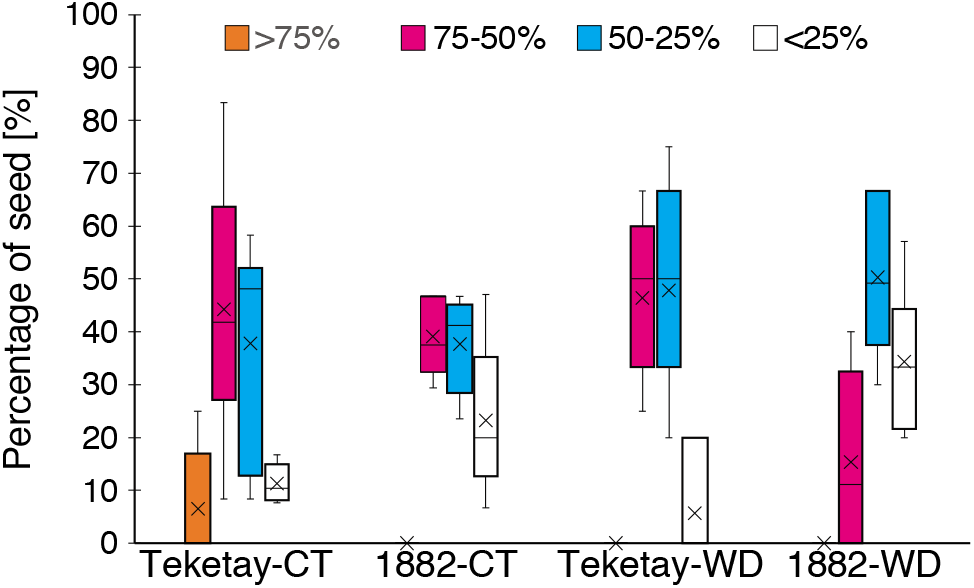
Distribution of seed development index of the chickpea genotypes Teketay and ICC 1882 in control (CT) and water deficit (WD) conditions. The box plots show the percentage of seeds distributed in each class of seed development index. The bottom, middle and top lines of the box represents the lower quartile, the median and the upper quartile, respectively. The median was excluded from quartile calculation when the number of replicates was odd. The cross represents the mean and the whiskers extend to the minimum and maximum values. Seeds were collected from five to seven plants.

**Table 2.**
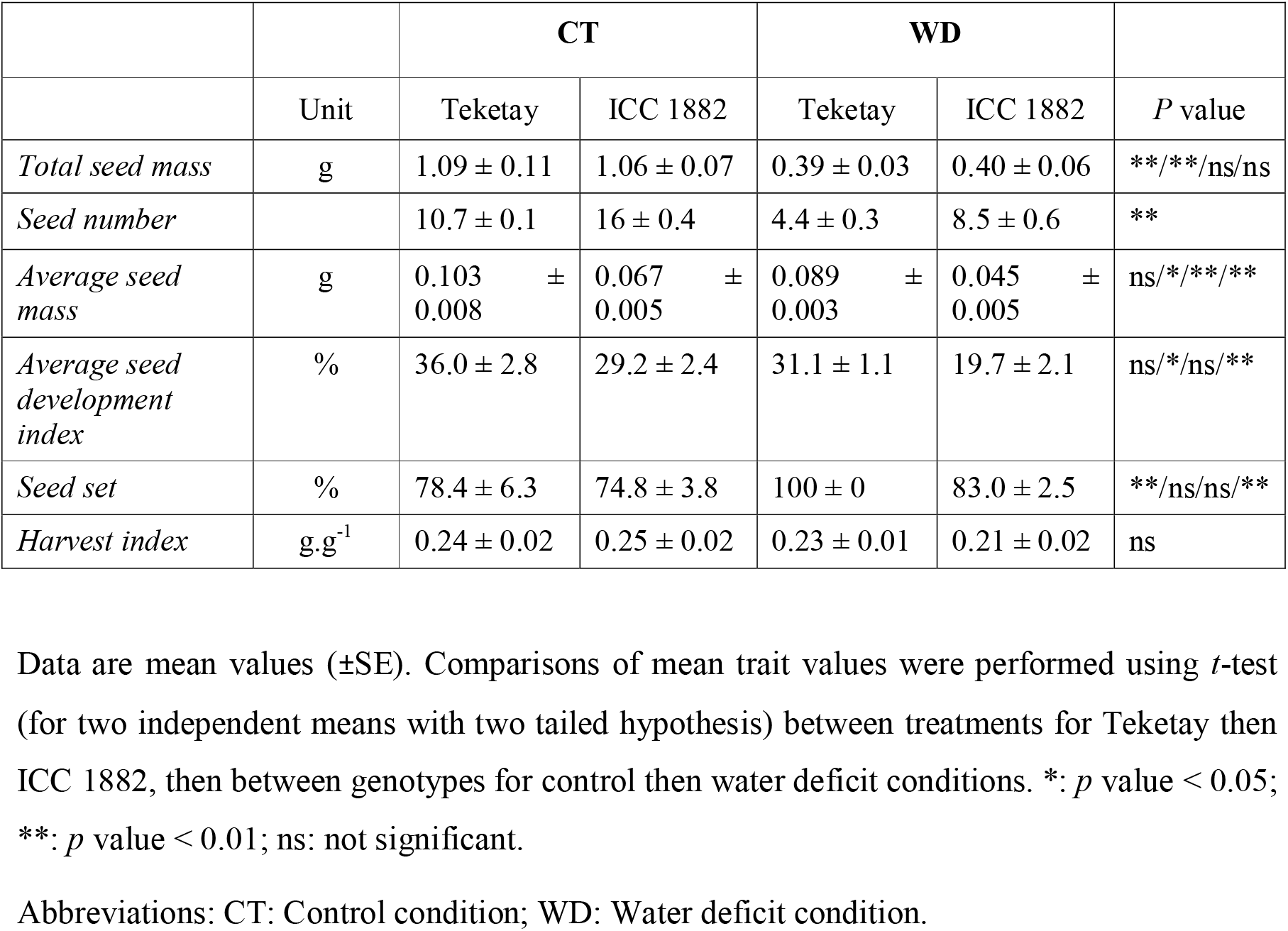
Seed related traits of the chickpea genotypes Teketay and ICC 1882 in control and water deficit conditions in rhizobox.

### Intrinsic differences between genotypes

Genotypes were also compared within treatments to highlight differences caused by genetic rather than environmental factors. We compared Teketay and ICC 1882 performance in CT then WD conditions. Overall, ICC 1882 shoot height was lower compared to Teketay in both conditions (Fig. 1). In CT conditions, it was only significant at the end of the vegetative phase (17-27 DAS). In WD conditions, Teketay did not exhibit a significantly increased shoot height consistently during the vegetative phase (only at 20 das, *p* < 0.05), but did during most of the reproductive phase (*p* < 0.01 at 38, 41 and 45 das).

Significant differences for RSA parameters were mainly detected during early vegetative stages in both conditions due to early vigour of Teketay (Fig. 2): Its root length was significantly higher at 10-24 das in CT conditions, and at 6-20 das in WD conditions. In both conditions, CHA and root depth were not significantly different between the two genotypes except for the following cases: Teketay CHA was higher at 20 das in CT condition; Teketay root depth was higher at 13 DAS and 6-13 DAS in CT and WD conditions, respectively. As a consequence, its solidity was also higher: this genotype invested more resources into root biomass formation, likely enabling more intense resource acquisition during the early vegetative phase (*p* < 0.05 at 10, 17-20 das in CT conditions, at 10-17 das in WD conditions). Its RGR was also significantly higher at 10-20 das in CT conditions but only at 13 das in WD condition (Fig. 2E, F).

Interestingly, we observed an RGR slowdown in Teketay at 27 das in WD condition, whereas for ICC 1882, RGR kept on increasing (Fig. 2F). As a result, root solidity was higher for ICC 1882 from that point onwards. From 31 das (Fig. 2F, *p* < 0.05), Teketay RGR was lower than ICC 1882. This slowdown in root development in the late vegetative phase delayed the RGR peak in Teketay by few days and resulted in higher RGR at 45 das (*p* < 0.05).

### Teketay root system development adapts to soil moisture and influences water use pattern

The differences in RA between genotypes, stratified by soil depth (ΔRA), were analysed during the vegetative (Fig. 8A, B) and reproductive phases (Fig. 8C, D). This analysis revealed an inflection zone (the stratum between 45-60 cm), below which root growth behaviour differed from the shallower layers.

**Figure 8.**
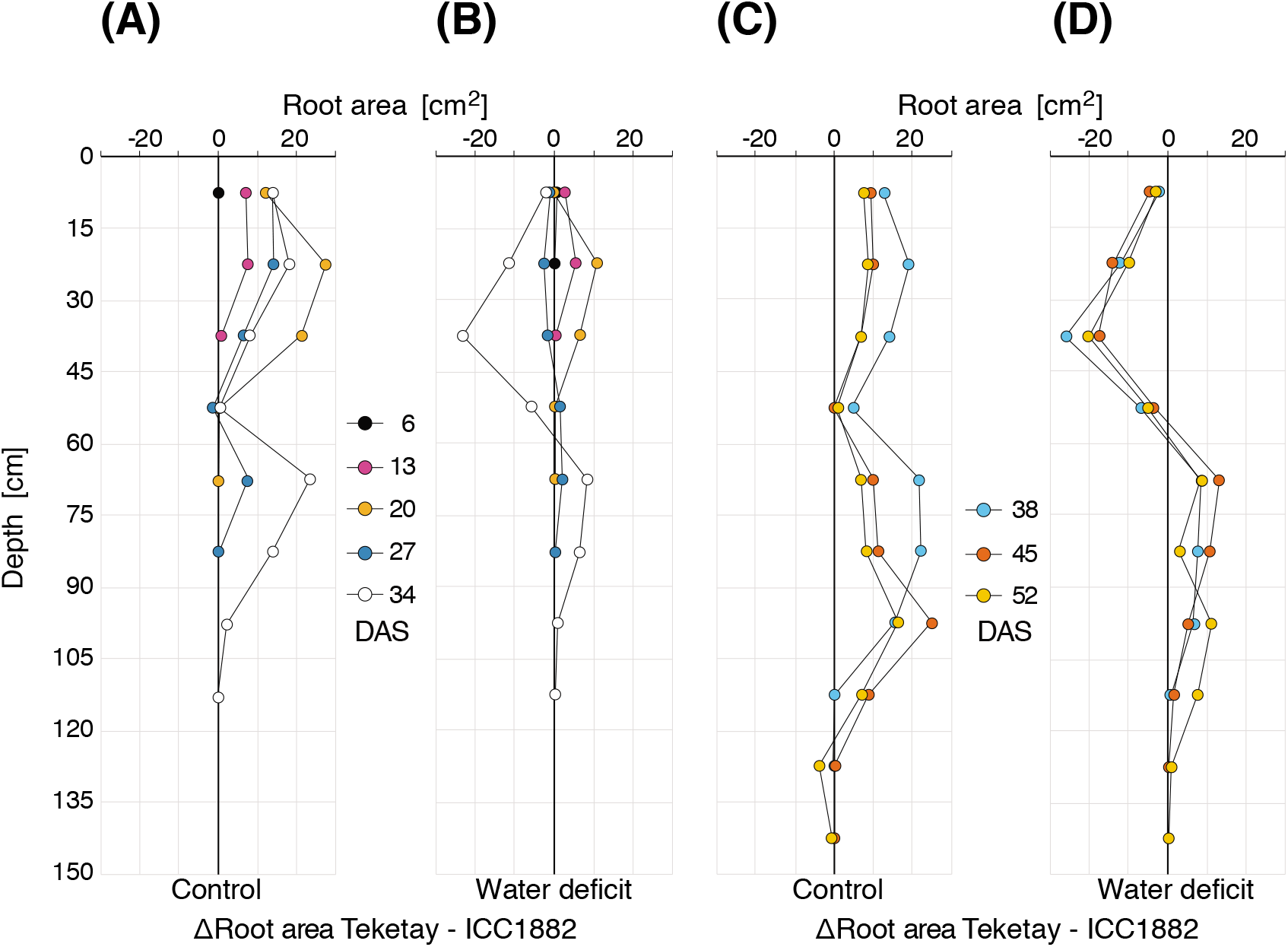
Spatio-temporal differences in root area (ΔRA) between the chickpea genotypes Teketay and ICC 1882 in control (CT) and water deficit (WD) conditions. ΔRA was calculated in 15 cm deep horizontal soil layers as RA_Teketay_ – RA_ICC_ 1882 in CT conditions during the vegetative (a) and reproductive (c) phases; in WD condition during the vegetative (b) and reproductive (d) phases. The legend indicates the number of days after sowing. To simplify the comparison between genotypes, the flowering time which marks the transition between vegetative and reproductive phases was set as 34 days after sowing as it was the closest time point to real flowering time for all conditions, except for Teketay which flowered at ~31 days after sowing in control conditions.

In CT conditions, Teketay clearly shows an increased RA compared to ICC 1882 for all soil strata, except at the latest stages (48-52 das) where marginally more RA was observed for the latter in some layers (45-60 cm and 120-150 cm depth, Fig. 8C). Steady ICC 1882 root growth progressively reduces the advantage Teketay acquired earlier: Within the same soil layer, ΔRA decreased over time as ICC 1882 root growth was delayed compared to Teketay. For example, ΔRA in favour of Teketay was maximal at 20 das in 15-45 cm depth in CT condition (Fig. 8A), then decreased while ICC 1882 catches up root growth in those layers (Fig. 8C).

In WD conditions, ΔRA was positive in top layers until 20 das reflecting early vigour of Teketay root growth (Fig. 8B). That pattern progressively inverted at the end of the vegetative phase (27-34 das) as ΔRA became negative in top strata (0-60 cm) but positive in deeper strata (60-120 cm, Fig. 8B). That trend was stable during the reproductive phase reflecting contrasting root investment strategies: upper layers for ICC 1882, deeper layers for Teketay (Fig. 8D).

In both soil moisture conditions, no significant differences for total WU were detected between genotypes, although ICC 1882 used on average ~8% more water than Teketay in WD conditions (Table 1). Cumulative WU was significantly higher for Teketay at 24 and 27 das in CT conditions (*p* < 0.05, Fig. 3A). Its WUR was also significantly higher at 24 das (*p* < 0.01, Fig. 3B). By contrast, in WD conditions, cumulative WU and WUR, respectively, were significantly higher for ICC 1882 at 10, 13 and 20 das (Fig. 3A), and at 20 and 27 das (Fig. 3B). However, Teketay WUR was significantly higher later in the reproductive phase (at 45 DAS).

Teketay extracted more water from the uppermost soil layer in CT conditions (+ 3.3% ΔSWC at 7.5 cm depth). At 82.5 cm depth, Teketay water extraction (ΔSWC) was significantly higher (+ 7%, *p* < 0.05) than ICC 1882’s in WD conditions (Fig. 4A, B).

### Soil moisture conditions differentially alter resource allocation in chickpea genotypes

Overall, whole plant and individual organ biomass, as well as relative biomass distribution, were similar for the two tested genotypes when grown in the same soil moisture condition (Table 1). However, we observed significant differences between genotypes in the following conditions: in CT conditions, stem DM and its ratio to plant DM were higher for Teketay compared to ICC 1882 (*p* < 0.01 for both, Table 1). By contrast, in WD conditions, the relative root DM was higher for Teketay (*p* < 0.05), indicating a higher investment into root biomass. In WD conditions the fraction of total DM allocated to leaves (*p* < 0.1) as well as the root/shoot DM ratio (*p* < 0.1) were markedly, although not significantly, lower and higher for Teketay, respectively. The harvest index was similar in both conditions (Table 2).

In CT conditions, Teketay flowered on average 2 days earlier than ICC 1882 (*p* < 0.05), but not in WD conditions (Table 1). In CT conditions, the number of reproductive organs was almost identical for both genotypes (Fig. 5A). In WD conditions, it was significantly lower for Teketay from 41 das. The numbers of pods as well as the pod set percentage were significantly lower for Teketay at 41 and 52 das in CT conditions (Fig. 5B, C). In WD conditions, the number of pods produced by Teketay was significantly lower from 41 das compared to ICC 1882, but their pod set percentage was very similar over time. Teketay pod number per root length was significantly lower than ICC 1882 at 41 then 48 and 52 das in CT condition and from 41 das in WD conditions (Fig. 6A). As a result, WUR per pod was significantly higher for Teketay at 41 das in CT conditions and from 45 das in WD condition (Fig. 6B).

Total seed mass was similar between the genotypes in both soil moisture conditions (Table 2). In both conditions, the number of seeds was significantly higher for ICC 1882 (*p* < 0.01). The percentage of seed set was similar between the genotypes in CT, but significantly higher for Teketay in WD conditions (*p* < 0.01). The average seed mass was higher (almost twice) for Teketay in both conditions, but the average seed development index was significantly higher for Teketay in WD conditions only (Table 2). The percentage of seeds in the development index classes < 25% and 50-75% was lower and higher, respectively, for Teketay in WD conditions (*p* < 0.01 for both, Fig. 7).

## Discussion

Drought is a major abiotic stress limiting plant development, reproduction and yield. Crops must adapt to minimize its impact (Tardieu *et al*., 2018). In conditions where annuals develop on residual soil moisture without irrigation, as is the case for chickpea cropped in rainfed agriculture, genotype-dependent root system plasticity and water management are critical to ensure reproduction and yield (Bodner *et al*., 2015). Our study utilises a non-invasive root phenotyping system (Bontpart *et al*., 2020) to compare the genotype-dependent responses to two levels of initial soil moisture. We show the importance of (*i*) RSA plasticity, (*ii*) RSA dynamics during vegetative and reproductive phases, and (*iii*) the coordination between RSA, water use and reproductive organ development for a successful drought resistance strategy.

### A large soil-based rhizobox system to dissect RSA responses to soil moisture during the crop life cycle

Soil-based rhizobox systems have previously highlighted significant variation between water regimes and genotypes at early developmental stages and were used as a pre-screening strategy to identify drought resistant varieties (*e.g*. Avramova *et al*., 2016). However, very few studies have reported WD effects over longer periods on plant development (Fang *et al*., 2017; Price *et al*., 2002). Due to root system growth after the vegetative phase (Bontpart *et al*., 2020), extending RSA phenotyping further into the crop life cycle is important to understand the plant’s water and growth management strategies.

We aimed to analyse the water budget of the chickpea plants in conjunction with RSA dynamics as comprehensively as possible. We grew chickpeas in rhizoboxes filled with two contrasting initial soil moisture levels and without further irrigation, to mimic the highly variable rainfall in (semi-) arid environments used for rainfed agriculture. The CT condition recapitulates a scenario where water is initially abundant and optimal for growth due to previous high rainfalls, but percolates by gravity to accumulate in deeper soil layers. This creates a soil moisture gradient which intensifies over time as upper soil strata become drier and deeper layers wetter, compared to initial soil moisture levels. Our WD conditions mimic a scenario where chickpea is sown after poor seasonal rains in a semi-arid environment and over its lifecycle is rapidly exposed to low soil moisture. In the latter conditions, soil moisture, albeit lower, is more homogeneous across the soil layers. Soil moisture decreases over time in both conditions from top to bottom due to gravity and progressive water extraction by the plant.

Overall, the magnitude of chickpea root traits (biomass, length, convex hull area, depth, ΔRA) was reduced in WD conditions. This is consistent with previous studies where WD decreased root length and CHA (Avramova *et al*., 2016; Durand *et al*., 2016), root biomass (Price *et al*., 2002) and root length density (Fang *et al*., 2017) in different plant species. Drought-induced root growth stimulation was only reported for Arabidopsis grown in rhizoboxes filled with a very thin soil layer (Rellan-Alvarez *et al*., 2015).

Field-grown chickpea root system dynamics have been evaluated using data obtained from only a fraction of the root system: from soil samples collected with coring tubes. In such experiments, the average root length density (per soil volume) of the chickpea genotype ICC 1882 was not clearly different between optimal irrigation and drought conditions, with distinct trends depending on developmental stage and season (Purushothaman *et al*., 2017b). However, a recent meta-analysis of field trials clearly demonstrates that plant (including legume) root length is reduced by drought, the degree of which depends on the specific drought scenario (Zhou *et al*., 2018).

### Teketay rapid development and lower pod set favoured better seed development in high initial soil moisture

The timing of the transition between vegetative and reproductive phases is critical for plant to ensure their reproduction (Jung and Muller, 2009). A key feature of annual plants adopting a drought *escape* strategy is early or accelerated flowering to complete reproduction before onset of severe drought (Berger *et al*., 2016; Grime, 1979). Such fast-growing chickpea genotypes have been successfully used by farmers to ensure yield in dry areas (Kumar and Abbo, 2001). When compared to ICC 1882, Teketay flowers almost 2 days earlier and completes pod set more rapidly in these conditions that mimic ‘normal’ rainfed agriculture.

Teketay exhibits an early vigour illustrated by higher above and belowground growth and more intense WU at the end of the vegetative phase (Table 1), which is consistent with an ‘escape’ strategy (Berger *et al*., 2016). By contrast, ICC 1882 growth is slower and delayed compared to Teketay; consistent with a previous study where ICC 1882 was described as a late vigour genotype (Sivasakthi *et al*., 2017). As a result, Teketay early vigour likely enables higher resource acquisition to reach the reproductive phase faster. However, the higher water consumption in upper soil layers that enables an accelerated transition from vegetative to reproductive phase would not be available during the reproductive phase. To compensate for its early high-consumption strategy in CT conditions, Teketay invested more resources into deep rooting (60-120 cm depth) during its reproductive phase for water extraction (Fig. 8).

Drought-resistant genotypes usually use more water during the reproductive phase compared to sensitive ones (Zaman-Allah *et al*., 2011b). However, WUR pattern, TR_mass_ during the reproductive phase, and ΔSWC in deep soil strata were not key factors for elevated drought resistance of Teketay compared to ICC 1882 in CT conditions. The most marked difference between the genotypes was distinct reproductive behaviour with Teketay limiting reproductive organ development, while ICC 1882 did not.

The number of reproductive organs produced by both genotypes was almost identical in CT conditions. However, not all flowers matured into pods, likely due to resource limitation. We observed that pods formed last exhibited limited development and produced no or very small seeds, consistent with a previous study (Leport *et al*., 2006). Teketay exhibited lower pod set and produced fewer pods than ICC 1882, but invested more resources into the development of each seed. As a result, each of its pods benefited from more water, leading to the production of further developed seeds (some reaching > 75% optimal seed biomass).

When resources allocated to reproduction are limited, a trade-off between seed size and number emerges in plants (Smith and Fretwell, 1974). Teketay seeds are 25% heavier than ICC 1882’s when plants are grown in optimal conditions; hence, it is likely that each Teketay pod requires more resources for optimal seed development. To compensate for bigger seed size, the Teketay reproductive strategy consisted of maturing a smaller fraction of flowers into pods (lower pod set) compared to ICC 1882, and as a result limiting the potential number of seeds. Seed number and maturity are key parameters for plant yield and therefore, the abortion of reproductive organs appears as beneficial trait during water limitation (Tardieu, 2012). In conclusion, Teketay escapes drought by rapid early development when water is still relatively abundant in high initial soil moisture, which allows for enhanced completion of its reproductive phase, albeit of fewer seed.

### Teketay late root investment in deep soil layers and lower pod production ensured better seed development in low soil moisture

Teketay responses to low initial soil moisture revealed a different strategy: they were not based on a shorter vegetative phase (‘drought escape’) compared to ICC 1882, as the two genotypes exhibit very similar FT, but founded on different RSA dynamics (Fig. 2F): Teketay developed a longer root system until mid-vegetative phase (Fig. 2A), although its shoot growth was similar to ICC 1882. The subsequently diminished Teketay RGR highlighted a change in its resource acquisition strategy, contrasting with its strategy of high consumption in CT conditions. Indeed, one trait (here WUR) can be beneficial for reproduction in a given drought scenario, but be detrimental in another one (Tardieu, 2012). Hence, Teketay reduced vegetative RGR and WUR to save water later used for reproduction, thereby revealing a drought postponement strategy.

In both growth conditions, Teketay pod number reached a plateau earlier than in ICC 1882. However, instead of aborting flowers like in CT conditions, it produced fewer reproductive parts but maintained a high percentage of pod set. In the meantime, its root growth increased in deep soil strata where its root system extracted more water as revealed by higher WU and TR. This observation fits with the report that drought tolerant genotypes exhibit high TR during the reproductive phase (Pang *et al*., 2017a).

In the field, water extraction in deeper soil layers (90-120 cm) is critical for drought adaptation since moisture decreases in higher soil layers as water evaporates from the surface and crops utilise it (Purushothaman *et al*., 2017a). Teketay resource investment into root growth in deeper soil strata was efficiently converted into higher WUR during the reproductive phase. Furthermore, in WD conditions, this genotype allocated more resources towards root biomass at the expense of stems when compared to CT conditions. This impact of WD on the root/stem biomass ratio is common in vascular plants as they invest relatively more into root rather than stem biomass when water availability is limited (Poorter *et al*., 2012).

We also observed a clear shift in Teketay shoot system architecture in WD conditions which only consisted of the main stem without branches, in marked contrast to its highly-branched shoot architecture in CT conditions (data not shown). This is consistent with a strategy to reduce leaf area, and hence water loss, in WD conditions. Furthermore, pod abortion is higher in secondary compared to primary branches (Fang *et al*., 2010); therefore, restricting shoot architecture to a single branch avoids pod abortion and permits a higher proportion of flowers to develop and mature seed.

The tight temporal coupling between controlled pod set and an intensification of deep rooting facilitated resource acquisition which likely enables Teketay seed development to directly benefit from the extracted resources: each individual pod has a higher specific root length and WU to service it. Consequently, the Teketay average seed development index was higher than in the ICC 1882 genotype, and strikingly, it was not significantly lower than in high initial soil moisture (Table 2).

## Conclusion

Using a root phenotyping system, we analysed the performance of two chickpea genotypes in distinct soil moisture conditions which allowed us to relate RSA and WU to shoot growth and reproduction.

The analysis of RSA development over time in conjunction with physiological parameters provides a new approach to elucidate robust strategies to counter water deficit and incorporate these into crop improvement efforts. Our work reveals that RSA plasticity and dynamic changes in growth (‘drought escape’ and ‘drought postponement’) and resource acquisition strategies, combined with exquisite control of reproduction, as observed in Teketay, are desirable traits for yield resilience.

This work reveals that crop response strategies to very low levels of soil moisture (WD) conditions vary dynamically in space and time. In a context where climate change is predicted to result in drier environments that restrict root growth, introduction of suitable spatiotemporal patterns of root development will be critical to ensure robust yields in less predictable environments.

## Supporting information

All supplemental information

## Abbreviations

*A*: Net photosynthetic CO_2_ assimilation rate per leaf area
CHA: Convex hull area
CT: Control condition
das: Days after sowing
DM: Dry mass
FM: Fresh mass
FT: Flowering time
*g*_s_: Stomatal conductance per leaf area
RA: Root area
RGR: Root growth rate
RSA: Root system architecture
R/S: Root/shoot ratio
SWC: Soil water content
TR_mass_: Transpiration rate per leaf mass unit
TWU: Total water use
VWC: Volumetric water content
WD: Water deficit
WU: Water use
WUE: Water use efficiency
WUR: Water use rate
ΔRA: Difference in root area
ΔSWC: Difference in soil water content

## Supplementary data

Supplementary data are available at *JXB* online.

*Figure S1*. Environmental conditions in the greenhouse.

*Figure S2*. Leaf gas exchange measurements on chickpea genotype ICC 1882 grown in control and water deficit conditions in rhizobox.

*Figure S3*. Gravimetric soil water content (SWC) at different depths in unplanted, soil-only rhizoboxes.

*Figure S4*. Segmented chickpea root system in rhizoboxes.

*Figure S5*. Validation of root analysis algorithm.

*Figure S6*. Spatio-temporal differences in root area (ΔRA) between initial soil moisture conditions in 15 cm deep horizontal soil layers.

*Figure S7*. Progression in root area change stratified by depth in 15 cm horizontal soil segments.

## Acknowledgments

We thank Sofia Hanson and Araliya Arnott for their help during rhizobox experiments. The research was funded by Global Challenges Research Fund (GCRF)-BBSRC (reference: BB/P023487/1). This research was carried out with resources provided by the Edinburgh Plant Growth Facility, a specialist service provider for plant growth at the University of Edinburgh.

## Author contribution

Conceptualisation: TB and PD; Methodology: TB, IR, VG, CC, SAT and PD; Software: VG and SAT; Validation: TB; Formal Analysis: TB; Investigation: TB, IR, CC and LCTS; Resources: AJM, AF, SAT and PD; Data curation: TB, VG, SAT and PD; Writing – original draft: TB; Writing – Review & Editing: TB, IR, VG, CC, LCTS, AJM, PD; Visualisation: TB; Supervision: TB and PD; Project Administration: TB; Funding Acquisition: AF, SAT and PD.

## Data availability statement

The data supporting the findings of this study are available from the corresponding author, upon request.

