## Supplementary material for "Chickpea (*Cicer arietinum* L.) root system architecture adaptation to initial soil moisture improves seed development in dry-down conditions": All supplemental information

### Supporting Information

***Suitability of rhizobox system for the study of plant responses to water deficit***

Soil water content (SWC) of unplanted rhizoboxes was determined at the mid-point (26 DAS equivalent) and end of the experiment (52 DAS equivalent) (Fig. **S3**). Gravity creates a SWC gradient in rhizoboxes; however, this gradient forms faster, and is steeper, in control (CT) conditions. A clear SWC gradient was already observed in control conditions at 26 DAS, whereas the soil moisture profile for WD conditions remains similar to the profile at the experiment's onset. At the experiment's end, a soil moisture gradient of ~17% is generated for CT conditions, where the top of the rhizobox is drier (50% SWC at 7.5 cm depth) compared to the bottom (67% SWC at 135 cm depth). By contrast, in WD conditions, the soil moisture gradient is only 6.5%, with all soil layers drier than in CT conditions (uppermost layer: 36.8% SWC; deepest layer: 43.3% SWC) (Fig. **S3**).

Our large rhizobox system was designed to monitor the majority of root system growth dynamically (Fig. **S4**). To evaluate how well the root system traits measured by this system correlate with the ground truth of total root mass, we tested the correlation between total root biomass after washing and length/area calculated by the algorithm used in the system described by Bontpart and colleagues at harvest time (53 DAS, Fig. **S5**). The length of the root system is well correlated with both root fresh mass (FM, $r^2$= 0.906, Fig. **S5a**) and dry mass (DM $r^2$= 0.805, **S5b**), and similarly, RA is well correlated with total root FM ($r^2$= 0.888, Fig. **S5c**) and DM ($r^2$= 0.793, Fig. **S5d**). Overall, we therefore considered the length and area of roots calculated by the algorithm to be good proxies for root biomass.

***Supplemental Figure legends***

**Figure S1**. **Environmental conditions in the greenhouse**. Air temperature (a) and relative humidity (b) recorded at canopy level. The bottom, middle and top lines of the box represents the lower quartile, the median and the upper quartile, respectively. The cross represents the mean and the whiskers extend to the minimum and maximum values excluding outlier points. The circles represent data considered as outliers.

**Figure S2**. **Leaf gas exchange measurements on chickpea genotype ICC 1882 grown in control and water deficit conditions in rhizobox**. Net photosynthetic $CO_2$ assimilation rate per leaf area ($A$, a) and stomatal conductance per leaf area ($g_s$, b) were measured on chickpea genotype ICC 1882 grown in control (CT) and water deficit (WD) conditions at first flower open. Data are mean values (±SE) of three plants. Comparisons of mean trait values were performed using *t*-test (for two independent means with two tailed hypothesis) between soil moisture treatments for ICC 1882. *: $p$ value $< 0.05$.

**Figure S3**. **Gravimetric soil water content (SWC) at different depths in unplanted, soil-only rhizoboxes**. For control (CT) and water deficit (WD) conditions, soil samples were harvested at the middle (26 DAS, n=1) and at the end (52 DAS, n=2) of the experiment to determine SWC. Data are mean values (±SE).

**Figure S4**. **Segmented chickpea root system in rhizoboxes**. Each image panel is the result of merging five independent Raspberry Pi camera images covering the whole rhizobox surface *via* customized algorithms. The examples are of a Teketay root system grown in high (a) and low (b) initial soil moisture, and an ICC 1882 root system developed in high (c) and low (b) initial soil moisture. All images are at the same scale. The scale bar represents 10 cm.

**Figure S5**. **Validation of root analysis algorithm**. For all rhizoboxes included in the study, chickpea roots were harvested at 53 days after sowing, washed and weighed. The panels show the correlation between root fresh mass and root length (a), root dry mass and root length (b), root fresh mass and root area (c), root dry mass and root area (d). The coefficient of correlation ($r^2$) is indicated for each linear regression.

**Figure S6**. **Spatio-temporal differences in root area (ΔRA) between initial soil moisture conditions in 15 cm deep horizontal soil layers**. ΔRA was calculated for the chickpea genotypes Teketay (a) and ICC 1882 (b) by subtracting root area for the water deficit (WD) conditions from root area for the control (CT) conditions. The legend indicates the number of days after sowing.

**Figure S7**. **Progression in root area change stratified by depth in 15 cm horizontal soil segments**. Root area is shown in control (CT) conditions for the chickpea genotypes Teketay

(a) and ICC 1882 (b), and in water deficit (WD) conditions for Teketay (c) and ICC 1882 (d). The legend indicates the number of days after sowing.

### Supporting Figures

Fig. S1.

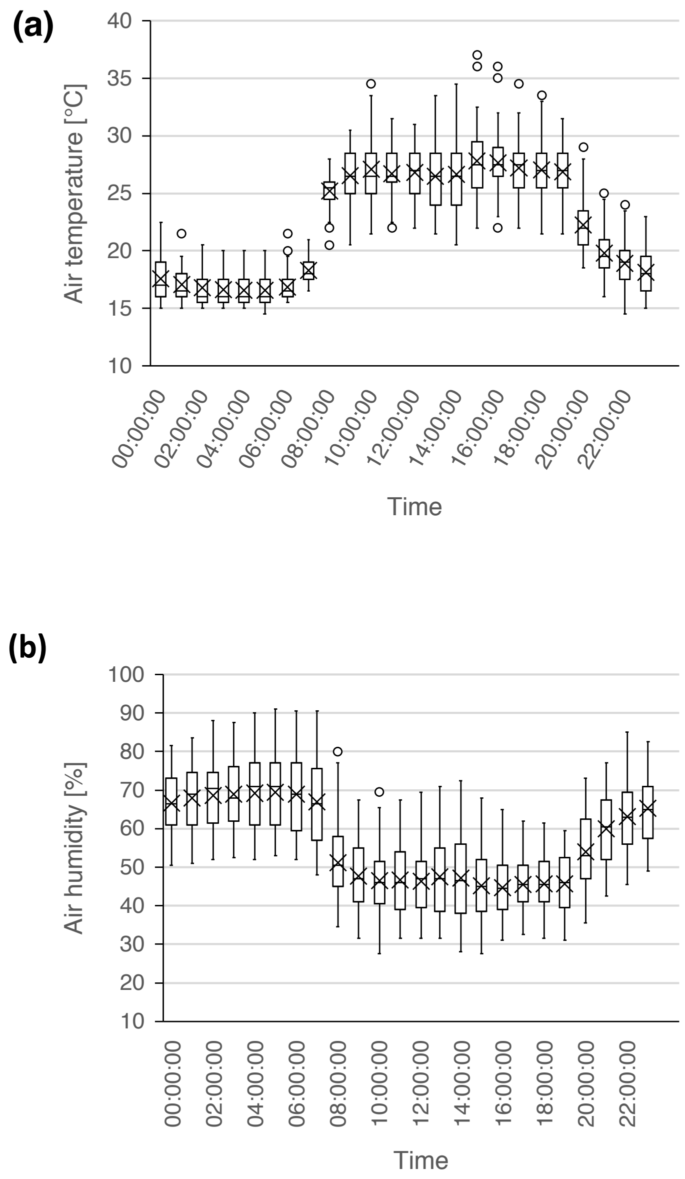


**Figure S1**. **Environmental conditions in the greenhouse**. Air temperature (a) and relative humidity (b) recorded at canopy level. The bottom, middle and top lines of the box represents the lower quartile, the median and the upper quartile, respectively. The cross represents the mean and the whiskers extend to the minimum and maximum values excluding outlier points. The circles represent data considered as outliers.

Fig. S2.

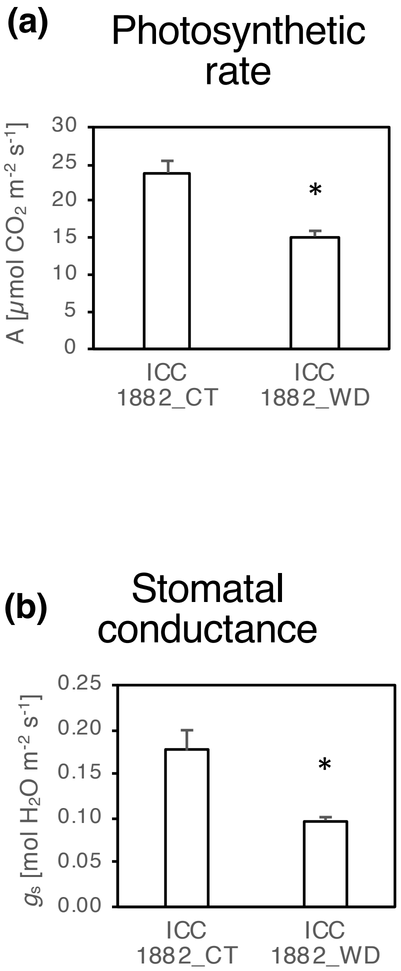


**Figure S2**. **Leaf gas exchange measurements on chickpea genotype ICC 1882 grown in control and water deficit conditions in rhizobox**. Net photosynthetic $CO_2$ assimilation rate per leaf area (*A*, a) and stomatal conductance per leaf area ($g_s$, b) were measured on chickpea genotype ICC 1882 grown in control (CT) and water deficit (WD) conditions at first flower open. Data are mean values (±SE) of three plants. Comparisons of mean trait values were performed using *t*-test (for two independent means with two tailed hypothesis) between soil moisture treatments for ICC 1882. *: $p$ value < 0.05.

Fig. S3.

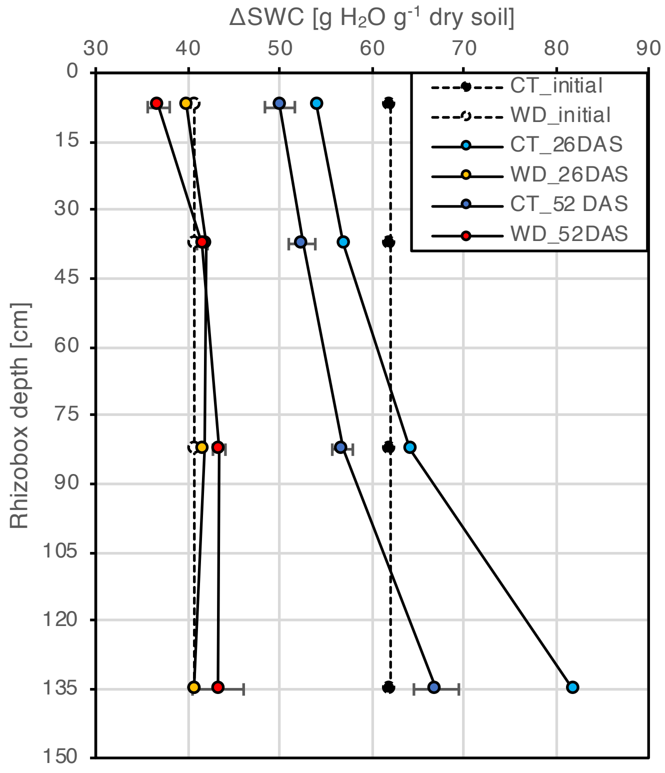


**Figure S3**. **Gravimetric soil water content (SWC) at different depths in unplanted, soil-only rhizoboxes**. For control (CT) and water deficit (WD) conditions, soil samples were harvested at the middle (26 DAS, n=1) and at the end (52 DAS, n=2) of the experiment to determine SWC. Data are mean values (±SE).

Fig. S4.

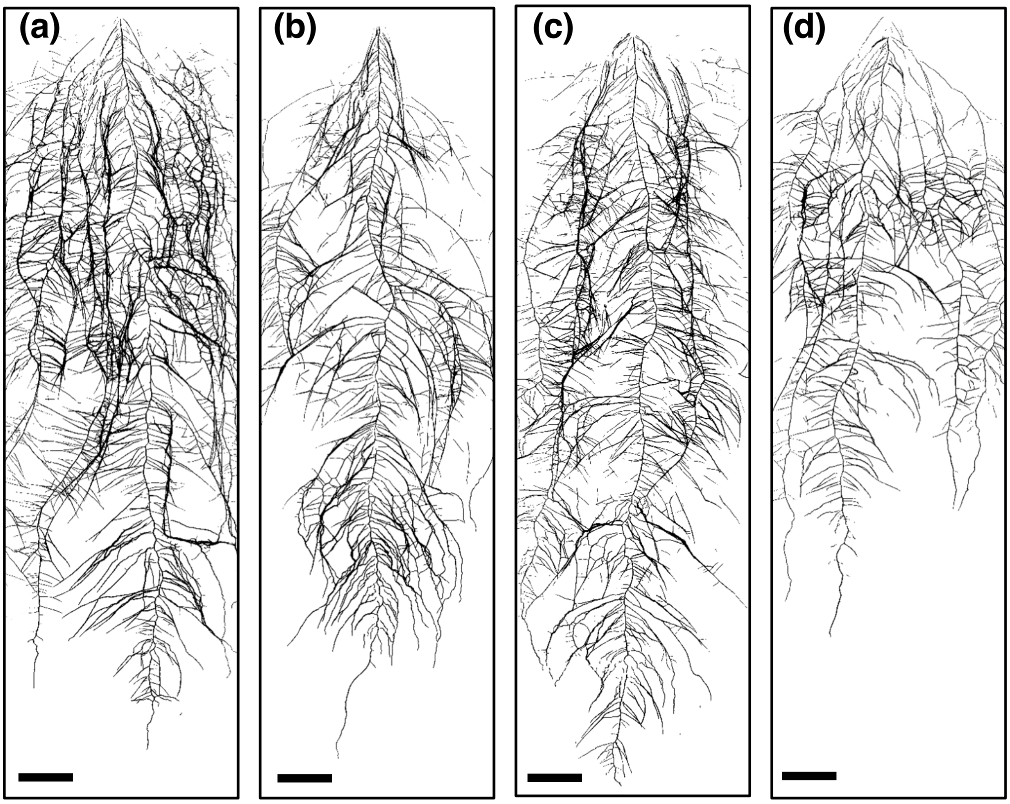


**Figure S4**. **Segmented chickpea root system in rhizoboxes**. Each image panel is the result of merging five independent Raspberry Pi camera images covering the whole rhizobox surface *via* customized algorithms. The examples are of a Teketay root system grown in high (a) and low (b) initial soil moisture, and an ICC 1882 root system developed in high (c) and low (b) initial soil moisture. All images are at the same scale. The scale bar represents 10 cm.

Fig. S5.

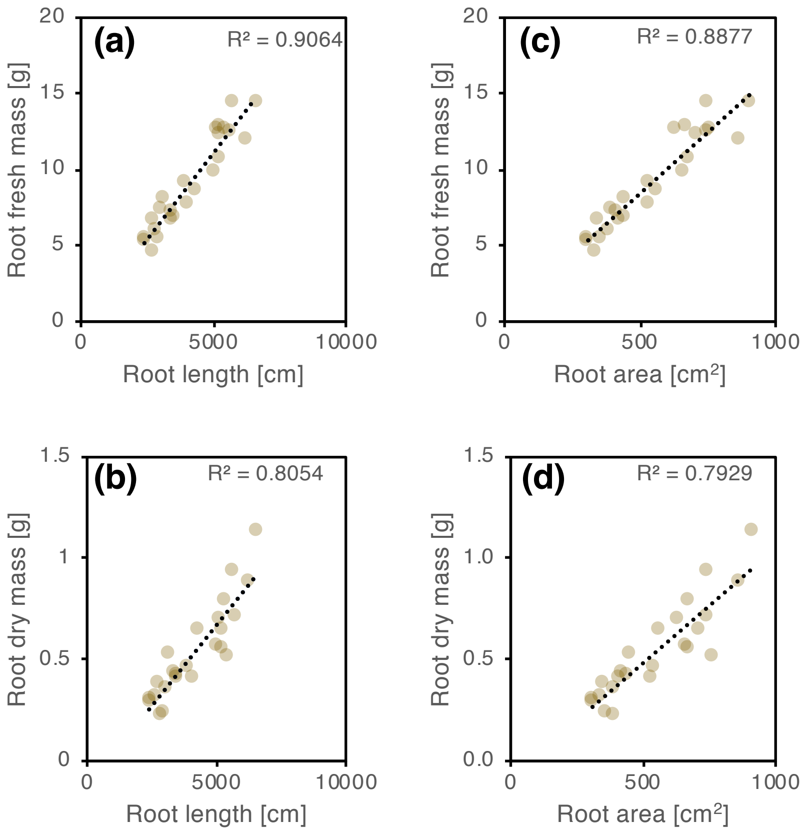


**Figure S5**. **Validation of root analysis algorithm**. For all rhizoboxes included in the study, chickpea roots were harvested at 53 days after sowing, washed and weighed. The panels show the correlation between root fresh mass and root length (a), root dry mass and root length (b), root fresh mass and root area (c), root dry mass and root area (d). The coefficient of correlation ($r^2$) is indicated for each linear regression.

Fig. S6.

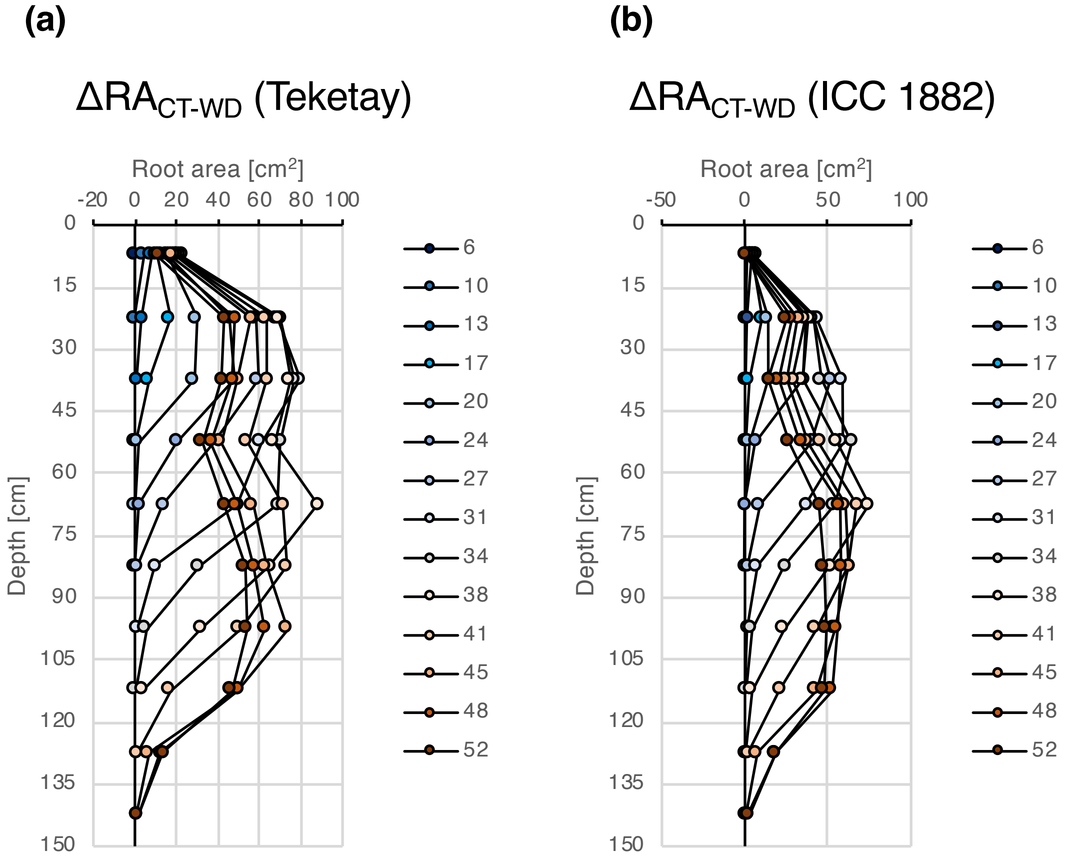


**Figure S6**. **Spatio-temporal differences in root area (ΔRA) between initial soil moisture conditions in 15 cm deep horizontal soil layers**. ΔRA was calculated for the chickpea genotypes Teketay (a) and ICC 1882 (b) by subtracting root area for the water deficit (WD) conditions from root area for the control (CT) conditions. The legend indicates the number of days after sowing.

Fig. S7.

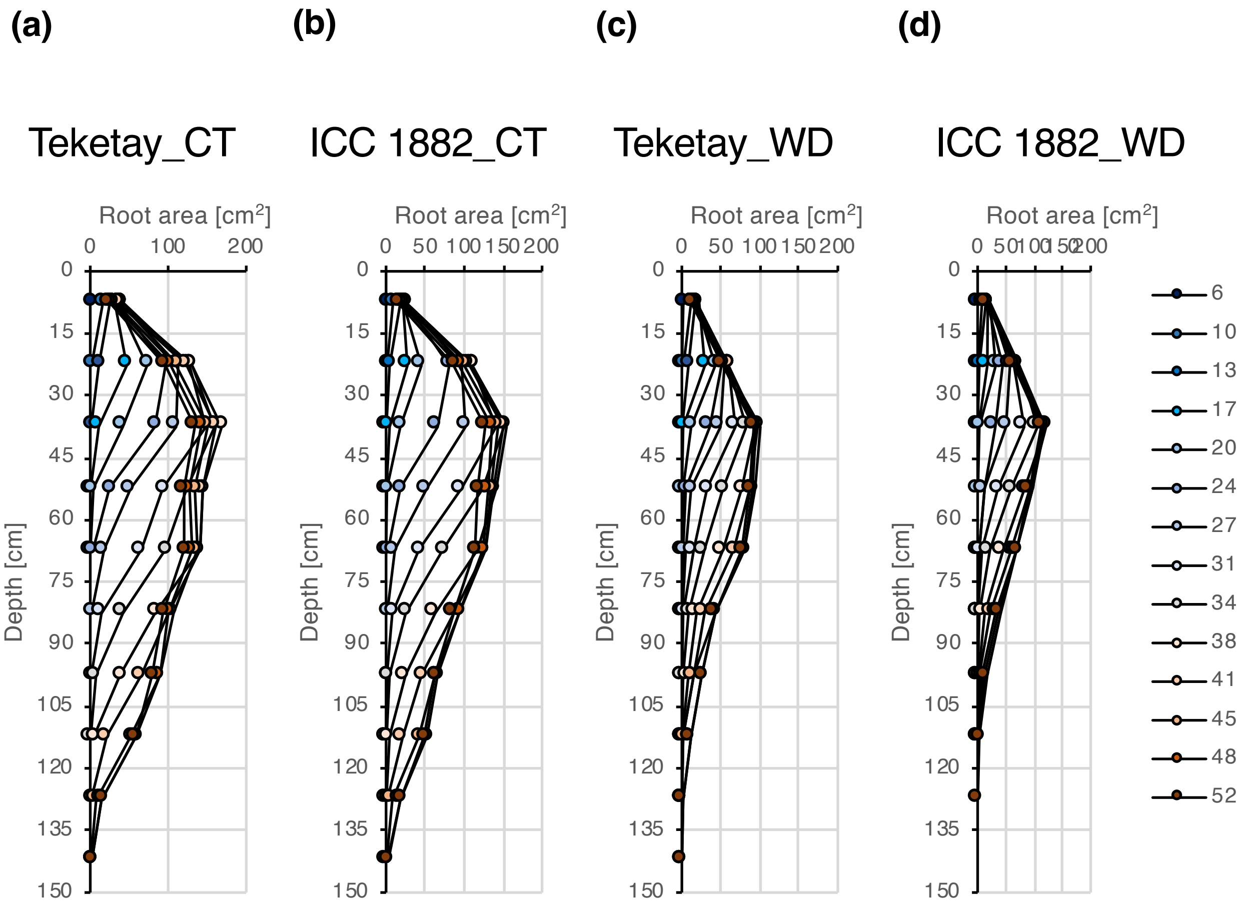


**Figure S7**. **Progression in root area change stratified by depth in 15 cm horizontal soil segments**. Root area is shown in control (CT) conditions for the chickpea genotypes Teketay (a) and ICC 1882 (b), and in water deficit (WD) conditions for Teketay (c) and ICC 1882 (d). The legend indicates the number of days after sowing.
